# Structure and Biosynthesis of Hectoramide B, a Linear Depsipeptide from the Marine Cyanobacterium *Moorena producens* JHB Discovered via Co-culture with *Candida albicans*

**DOI:** 10.1101/2023.07.06.547815

**Authors:** Thuan-Ethan Ngo, Andrew Ecker, Aurora Guild, Ariana Remmel, Paul D. Boudreau, Kelsey L. Alexander, C. Benjamin Naman, Evgenia Glukhov, Nicole E. Avalon, Vikram V. Shende, Lena Gerwick, William H. Gerwick

**Author notes:** Corresponding Author **William H. Gerwick –** Center for Marine Biotechnology and Biomedicine, Scripps Institution of Oceanography and Skaggs School of Pharmacy and Pharmaceutical Sciences, University of California San Diego, La Jolla, California 92093, United States.

## Abstract

The tropical marine cyanobacterium *Moorena producens* JHB is a prolific source of secondary metabolites with potential biomedical utility. Previous studies of this strain led to the discovery of several novel compounds such as the hectochlorins and jamaicamides; however, bioinformatic analyses of its genome suggested that there were many more cryptic biosynthetic gene clusters yet to be characterized. To potentially stimulate the production of novel compounds from this strain, it was co-cultured with *Candida albicans*. From this experiment, we observed the increased production of a new compound that we characterize here as hectoramide B. Bioinformatic analysis of the *M. producens* JHB genome enabled the identification of a putative biosynthetic gene cluster responsible for hectoramide B biosynthesis. This work demonstrates that co-culture competition experiments can be a valuable method to facilitate the discovery of novel natural products from cyanobacteria.

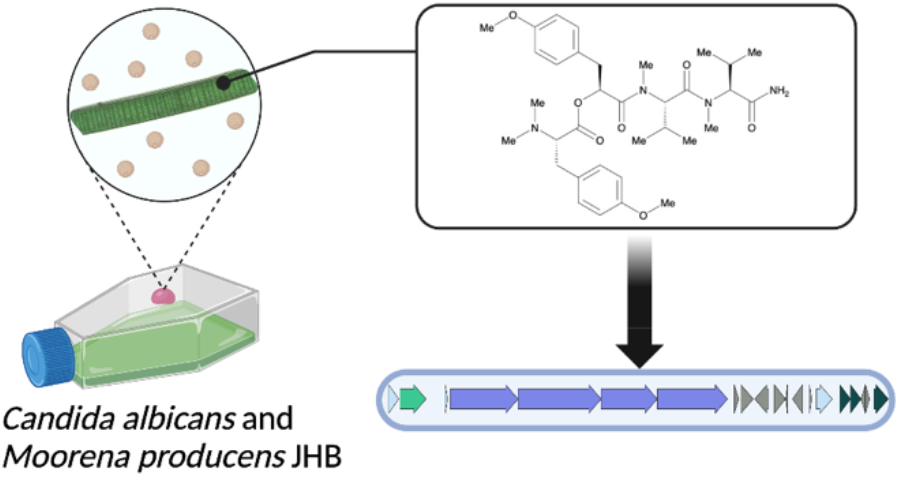

## Introduction

Members of the cyanobacterial genus *Moorena* are tropical, filamentous, photosynthetic, non-diazotrophic, and generally reside in the marine benthic zone.^1^ Notably, they are also a prolific source of bioactive secondary metabolites such as apratoxin A, a cyclic depsipeptide isolated from *Moorena bouillonii* collected in Guam that is a potent inhibitor of protein synthesis.^2–4^ Comparative genomics of species of this genus have revealed their extensive biosynthetic potential (∼18% of their genomes is dedicated to secondary metabolism) with the average number of biosynthetic gene clusters (BGCs) generally much higher than other cyanobacteria.^5^ Intriguingly, a significant number of these BGCs are silent or cryptic and have yet to be explored for their encoded products. Therefore, new approaches are needed to induce or increase the production levels of these natural products (NPs).

The tropical filamentous marine cyanobacterium *Moorena producens* JHB (formerly known as *Lyngbya majuscula*, hereinafter referred to as JHB) was obtained from a shallow marine environment in Hector’s Bay, Jamaica and has been continuously cultivated in seawater-BG11 (SWBG11) culture medium since its original collection.^6^ It is an abundant producer of diverse bioactive secondary metabolites, including the potent antifungal NPs hectochlorins A-D,^6^ sodium channel antagonists jamaicamides A-F,^7,8^ cryptomaldamide,^9^ and hectoramide A.^7^ Although this represents a large number of compounds isolated from a single cyanobacterium, genome analysis of this strain revealed that there are as many as 42 cryptic gene clusters yet to be investigated, 11 of which contain non-ribosomal peptide synthetase (NRPS)-related genes.^5^ Therefore, we sought to drive the expression of some of these previously uninvestigated BGCs through stimulation by a competing microorganism (*Candida albicans*), a method well known to upregulate the expression of NPs.^10,11^

## Results and Discussion

### Discovery from Co-culture Experiment

The biomass of the co- and mono-cultures of JHB with and without *C. albicans* were harvested, extracted, and analyzed, all in triplicate, by LCMS after 4 weeks of incubation. The peak areas of metabolites observed by LCMS were compared in order to identify metabolites with enhanced production in co-culture experiments as compared to the controls. We observed that the co-cultures had an antagonistic effect on the growth of the cyanobacteria, with biomass of co-cultured cyanobacteria greatly reduced compared to the mono-cultures after the same growth period. The production of hectoramide B (**1**) in the co-culture with JHB and *C. albicans* appeared increased relative to the mono-culture of JHB (Figure S14). This possible metabolic upregulation inspired interest in identifying and characterizing the structure, biosynthetic gene cluster, and bioactivity of this unique metabolite.

### Structure Determination of Hectoramide B (1)

Hectoramide B (**1**) was isolated by VLC and HPLC to yield 1.7 mg of yellow oil from the co-culture experiments. Based on HRMS, (exact mass = 627.3750 *m/z*), a putative molecular formula for compound **1** was calculated as C_34_H_50_N_4_O_7_. The 12 degrees of unsaturation required for this molecular formula were deduced by interpretation of NMR data to result from two phenyl rings and four ester/amide-type carbonyls.

The ^1^H NMR spectrum of compound **1** was remarkably similar to that of the previously determined structure of hectoramide^7^ [here renamed hectoramide A (**2**)]. Moreover, analysis of the MS/MS fragmentation using GNPS molecular networking,^12^ also revealed that **1** was closely related to **2**. However, its larger size suggested that it may have an additional amino acid residue in comparison with **2**. Additional NMR signals in **1** that were not present in **2** included aromatic proton peaks, *N*- and *O*-methyl singlets, and a deshielded proton alpha to a heteroatom coupled to a mid-field methylene group, altogether suggesting this additional amino acid might be an *N, N-*dimethyl-*O*-methyl tyrosine moiety.

SMART-NMR^13^ analysis of the HSQC spectrum of **1** further suggested the presence of multiple valine residues as well as methylated tyrosine residues (Figure S7). HSQC, HMBC and COSY correlations confirmed the sections of compound **1** that were identical to compound **2**, and also established the new residue as the proposed trimethyl-tyrosine residue (Figure 2a). This was also supported by analysis and comparison of the MS/MS fragmentation for **2** and **1**, which revealed some shared MS^2^ peaks (Figure 2b). Previous determination of the absolute configuration of **2**^7^, combined with a bioinformatic analysis as detailed below, were used to infer the absolute configuration of **1**. These analyses allowed deduction of the configurations of the tyrosine and both valine derivatives as L, and the Mppa moiety as *S*.

**Figure 1.**
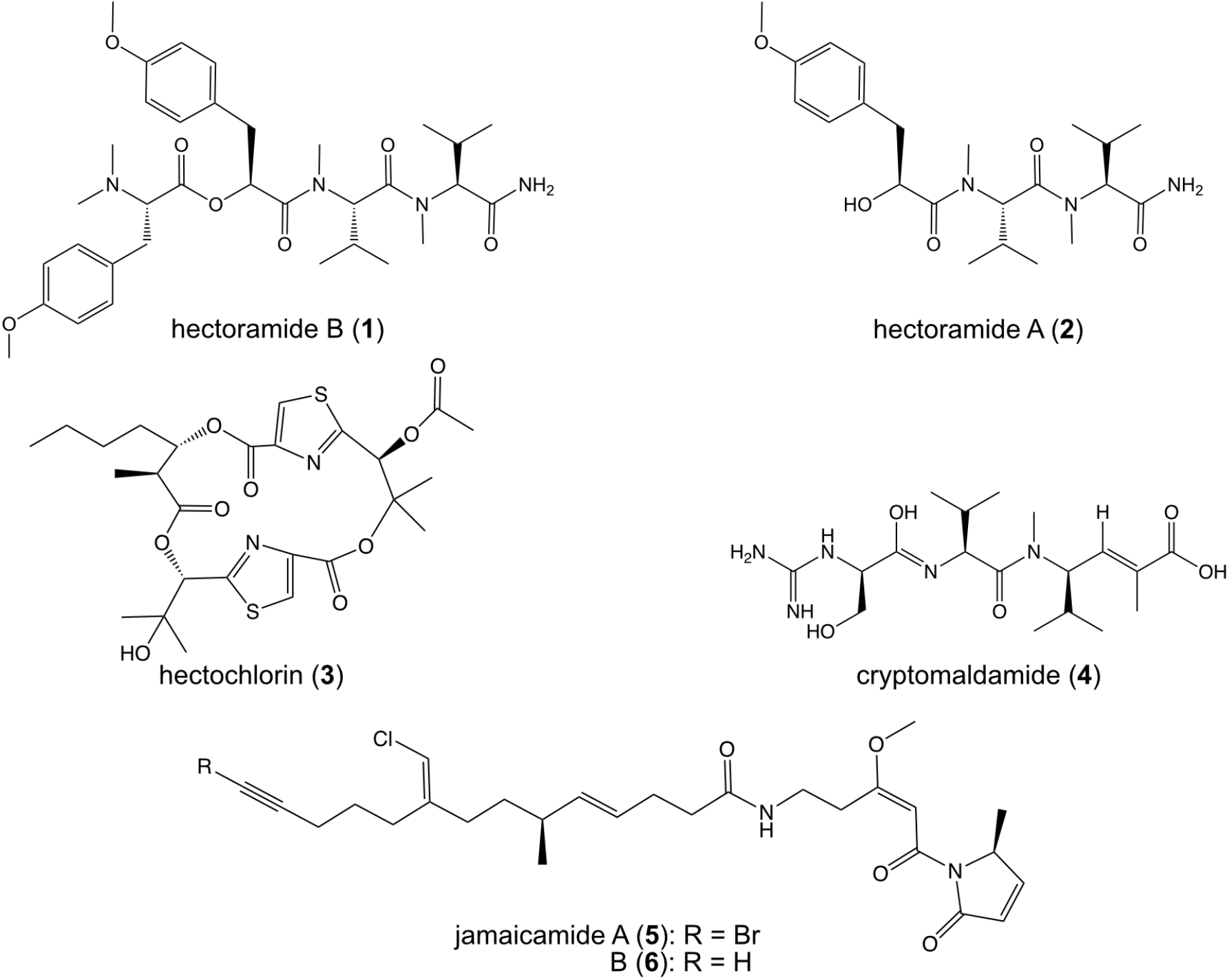
Natural products isolated and characterized from *Moorena producens* JHB

**Figure 2.**
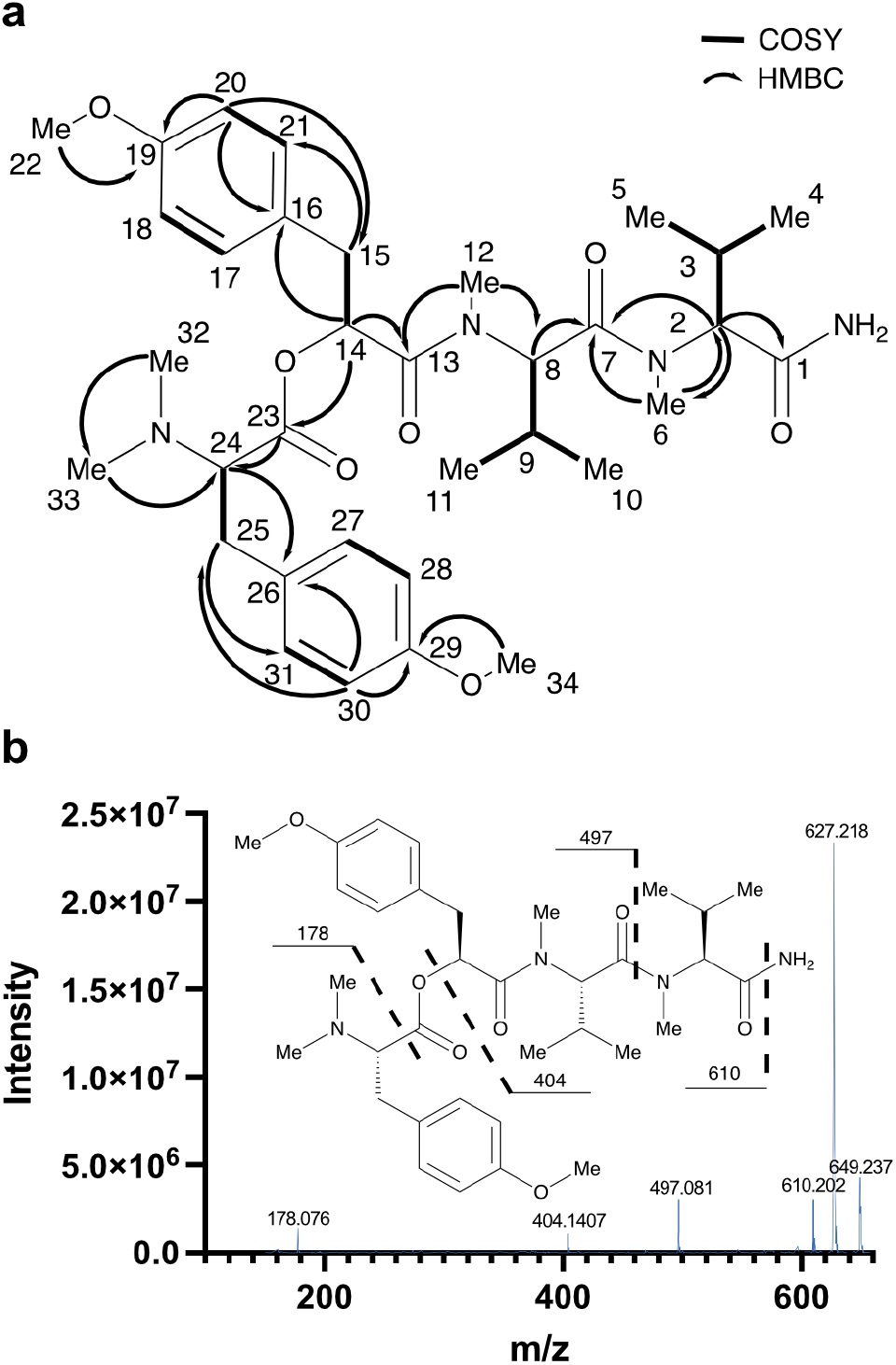
Structural assignment of hectoramide B. (**a**) Key COSY and HMBC correlations used in the structural determination of hectoramide B (**1**). All COSY and HMBC correlations are reported in Table S1. (**b**) Fragmentation pattern of **1** based on MS^2^ spectrum from the molecular ion (m/z 627.218)

### Biosynthetic Gene Cluster Analysis

Non-ribosomal peptides are typically synthesized in a collinear manner where each module encodes for the incorporation of an amino or hydroxy acid residue and chain elongation occurs in the sequence that the modules are ordered.^14^

A retrobiosynthetic scheme for the probable hectoramide B biosynthetic gene cluster was developed based on its chemical structure, including modules consistent with a non-ribosomal peptide synthetase (NRPS) incorporation of amino or hydroxy acids, and tailoring enzymes for key structural modifications such as methyl groups on heteroatoms (Figure S8). We hypothesized that the initial module would be an NRPS that would contain an adenylation (A) domain specific for tyrosine incorporation, followed by methyltransferase domains that would catalyze the methylations of the tyrosine phenolic oxygen atom and nitrogen atom of the amine group. We predicted that module 2 would include a depsipeptide synthetase that would incorporate an α-keto acid version of tyrosine that is subsequently reduced to 2-hydroxy-3-(4-hydroxyphenyl) propanoic acid by a ketoreductase (KR) domain. This module should also include a methyltransferase domain that methylates the phenolic oxygen atom of this tyrosine-derived residue. Modules 3 and 4 are predicted to contain A domains that incorporate valine residues followed by methylation by an *N*-methyltransferase. A terminal amidation enzyme is predicted to be encoded at the distal end of module 4, possibly related to one observed in carmabin A^10^ and vatiamide E and F^11^ biosynthesis, completing the pathway.

Previous sequencing efforts of the JHB genome were conducted using Illumina MiSeq and assembled using a *Moorena producens* PAL reference assembly.^5^ We were unable to identify a candidate biosynthetic gene cluster after initial inspection of this assembly. Therefore, we sought to obtain an improved genome assembly of JHB by extracting high molecular weight DNA and obtaining long-reads with Nanopore PromethION sequencing. Previous sequencing data from Illumina MiSeq and the new sequencing data from Nanopore PromethION were assembled with different tools to obtain an improved assembly of the JHB genome (Table 1). The final assembly of the JHB genome resulted in a single circular scaffold of 9.6 Mbp, a GC content of 43.67%, and completeness of 99.22%. Therefore, we selected this assembly for BGC analysis using antiSMASH (GenBank Accession Number: CP017708.2).

**Table 1.** Evaluation of the Quality of Genome Assembly Based on Quast and BUSCO^a^ analysis.

| Assembly method | # of Contigs | N50 | Total length (bp) | Complete<sup>a</sup> (%) |
| --- | --- | --- | --- | --- |
| SPades | 3 | 9373345 | 9384763 | 98.84 |
| Unicycler long-read | 2 | 8738093 | 9632771 | 60.67 |
| Unicycler hybrid | 2 | 5885695 | 9639251 | 99.22 |
| Flye | 1 | 9645659 | 9645659 | 79.56 |
| Flye + Pilon | 1 | 9648534 | 9648534 | 99.22 |

<sup>a</sup>Completeness was based on presence of complete single-copy orthologs in the BUSCO cyanobacteria\_odb10 database

### Putative Biosynthetic Gene Cluster for Hectoramide B

Inspection of the *M. producens* JHB genome revealed one candidate BGC that was consistent with the biosynthesis of hectoramide B (the *hca* pathway, MiBIG Accession: BGC0002754, GenBank Accession: OQ821997), the predicted retrobiosynthetic scheme, and the NMR and MS based structural assignment of hectoramide B (**1**) (Figure S8, Table S9). AntiSMASH annotations were integrated with protein family homology analysis, substrate selectivity predictions, and active site and motif identification. The *hca* pathway is composed of 4 core biosynthetic NRPS modules (*hca*A-*hca*D), flanked by putative regulatory genes, transport-related genes, and coding regions for hypothetical proteins, providing provisional boundaries to the biosynthetic cluster (Figure 3).

**Figure 3.**
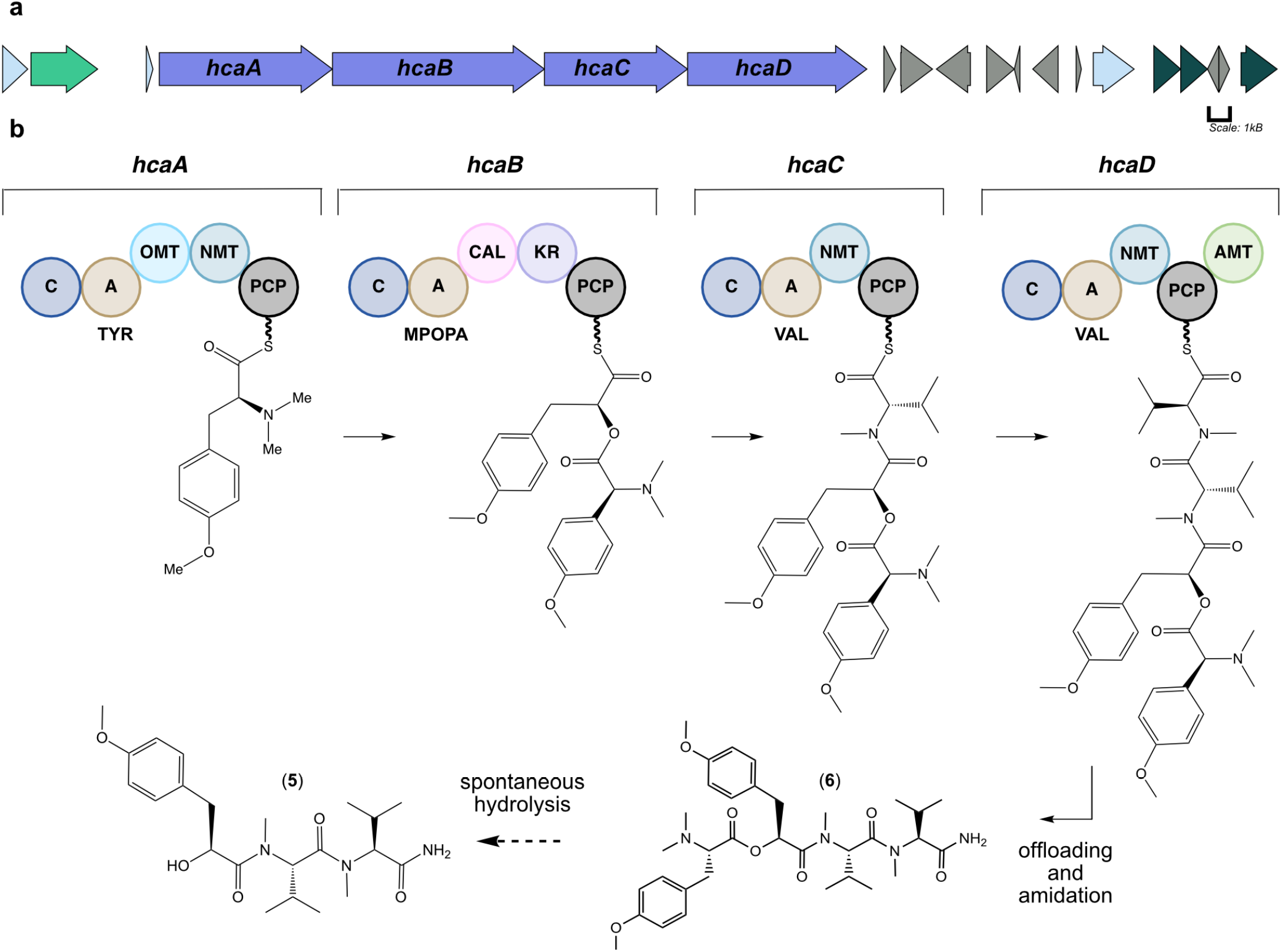
Putative biosynthetic gene cluster for hectoramide B. (a) Purple arrows indicate core biosynthetic genes. Additional arrows indicate additional ORFs that provide provisional boundaries to the hca BGC. Their proposed functions can be found in Table S9. (b) There are four core biosynthetic modules organized in a colinear fashion in the hca pathway. C: condensation domain, A: adenylation domain, TYR: adenylation domain for tyrosine incorporation, OMT: oxygen methyltransferase domain, NMT: nitrogen methyl transferase domain, MPOPA: adenylation domain for the proposed 3-(4-methoxyphenyl)-2-oxopropanoic acid incorporation, CAL: co-enzyme A ligase domain, KR: ketoreductase domain, VAL: adenylation domain for valine incorporation, AMT: amidotransferase. PCP: peptidyl-carrier protein. Phosphopantathiene arms are symbolized by wavy lines.

Bioinformatic analysis of the *hca* gene cluster suggested that the initial core biosynthetic module, *hcaA*, encodes for a multienzyme that is responsible for incorporating a tyrosine residue. This module contains condensation (C), adenylation (A), *O*-methyltransferase (OMT) and *N*-methyltransferase (NMT) domains as well as a peptidyl carrier protein (PCP) domain. The presence of both an OMT and the NMT is consistent with the amino terminus structure of hectoramide B. However, initial AntiSMASH analysis predicted the presence of two NMT domains, and therefore a phylogenetic tree of OMT and NMT domains from cyanobacteria was generated to examine more carefully the specificities of each of the annotated methyltransferase domains in *hcaA*. This revealed that the first MT, HcaA-MT1 clades well with other OMT domains, such as the OMT in the VatN module of the vatiamide biosynthetic gene cluster^15^ (Figure S10). The vatiamide OMT is also predicted to methylate the phenolic oxygen atom of a tyrosine residue. The second MT, HcaA-MT2, clades well with other NMT domains, specifically those within the *hca* pathway and the NMT of the VatN module in the vatiamide pathway. Therefore, *N, N-*dimethyl-*O*-methyl-L-tyrosine serves as the first structural unit in the *hca* pathway.

The second module, *hcaB*, is predicted to encode for a protein that ultimately incorporates 2-hydroxy-3-(4-methoxyphenyl) propanoic acid (Mppa). In previous studies of cyanobacterial depsipeptide formation, the A domain initially selects for an α-keto acid substrate that is reduced *in cis* to an α-hydroxy acid by a ketoreductase (KR) domain before incorporation into the natural product scaffold.^12^ However, antiSMASH analysis of this module did not reach a consensus for substrate specificity, and there was no annotation for an OMT domain to install a methyl group on the phenolic oxygen atom. Therefore, a sequence and structural alignment of the HcaB-A domain with other NRPS A domains selecting for tyrosine and phenylalanine was generated to identify the specificity conferring residues of hcaB-A (Figure 4a). Interestingly, the proposed specificity conferring residues of HcaB-A did not coincide with previous patterns observed in α-keto acid selecting A domains. Typically, in keto-acid activating A domains the conserved Asp235 residue, which traditionally hydrogen bonds with the primary amine of amino acids, is substituted to an aliphatic residue while the remaining specificity conferring residues match those predicted for the corresponding amino acid.^16^ However, neither was the case for HcaB-A; Asp235 is still present, and the remaining proposed specificity conferring residues did not match those expected for tyrosine (Figure 4a). Furthermore, alignment of a predicted structural model constructed *de novo* with AlphaFold2^17^ of HcaB-A and crystallography-derived structures of other NRPS A domains showed excellent congruence with the specificity conferring codes suggested by the sequence alignment described above (Figure 4b). Finally, as noted there was no annotated OMT domain in the *hcaB* gene. One possibility that is consistent with these features is that hcaB could be selecting for the α-keto acid form of *O*-methyl-tyrosine, 3-(4-methoxyphenyl)-2-oxopropanoic acid (MPOPA), rather than the α-keto acid form of tyrosine. An additional methyl group on the phenolic oxygen atom would require a significant alteration of the specificity binding pocket in order to accommodate this bulkier side chain.

**Figure 4.**
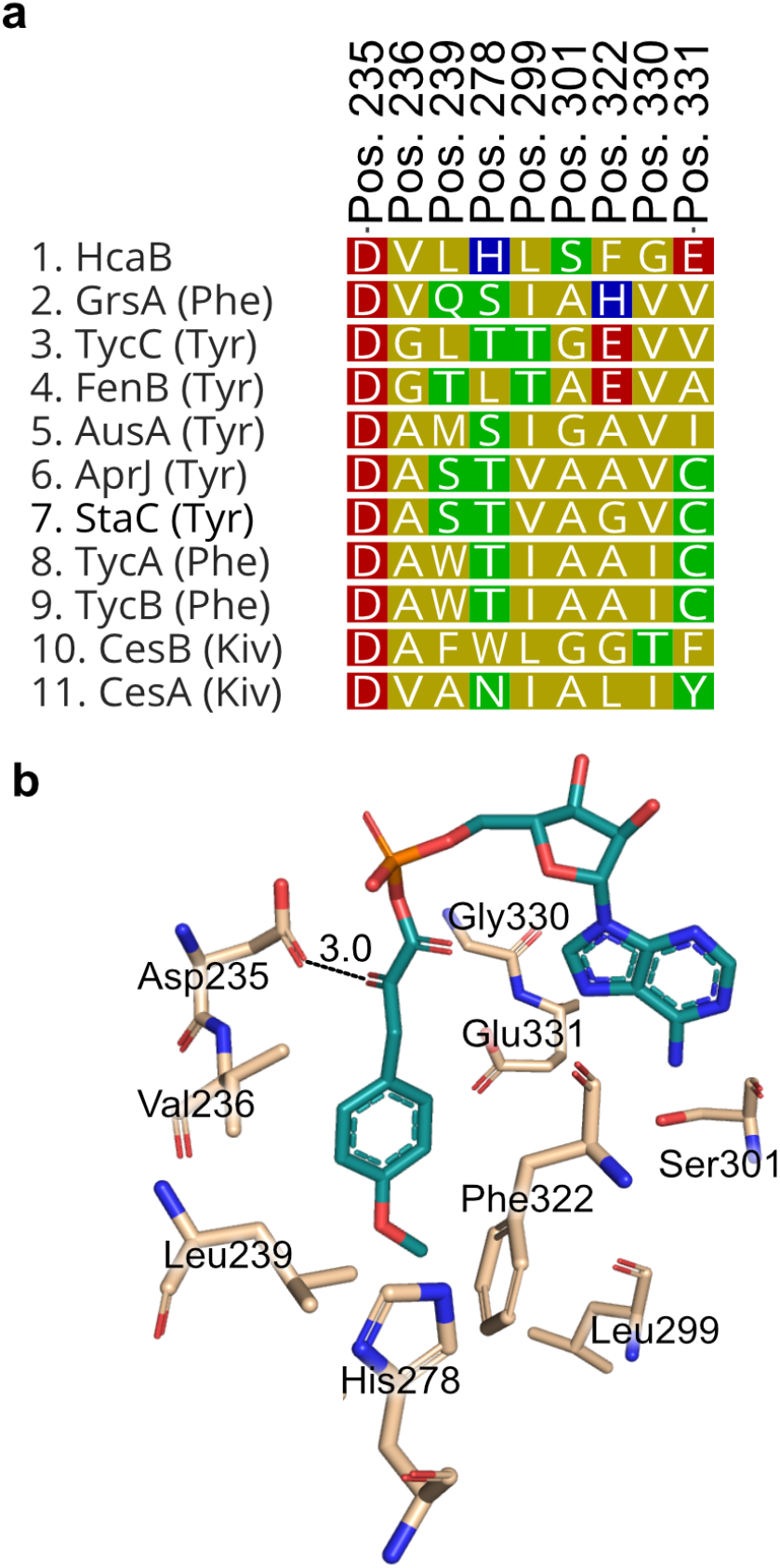
Sequence and structural alignment of hcaB adenylation domain reveals amino acids potentially conferring specificity, (a) Putative specificity conferring codes of HcaB-A compared with those from other adenylation domains from NRPS systems, (b) Three-dimensional model of hcaB-A domain generated using Alphafold2 bound to 3-(4-methoxyphenyl)-2-oxopropanoic acid (MPOPA) adenylate. The α-keto group of the Mpopa (teal) potentially binds through an antiparallel carbonyl-carbonyl interaction with the carbonyl group of Asp235 (wheat). See Table S12 for compound identities and other information

α-Keto acids can also be selected for by an antiparallel carbonyl-carbonyl interaction between the α-keto group and the carbonyl of the peptide bond connecting Gly414 and Met415.^18^ This carbonyl-carbonyl interaction has a comparable strength to that of a hydrogen bond and is important to α-keto acid selection. However, based on the model of HcaB-A (Figure 4b), the only residue that is in close proximity to the α-keto group is Asp235 (3.0 Å) whereas the Gly-Met backbone carbonyl is far distant. The predicted antiparallel orientation of these two carbonyls suggests a basis for stabilizing the incorporation of this α-keto acid.

Another component of this module is the domain annotated as a coenzyme A ligase (CAL). CAL domains are predicted to have specificity for fatty acids; however, as there are no fatty acid moieties in hectoramide B, it may be non-functional. Interestingly, similar CAL domains are found in a number of other depsipeptide producing BGCs such as cryptophycin-327,^19^ hectochlorin,^20^ and didemnin B,^21^ and therefore could be playing another role in the production of depsipeptides, a hypothesis that warrants further investigation. Lastly, the KR domain in the HcaB module is predicted to have stereospecificity for an *S* product. We propose that after activation of MPOPA, the KR domain in this module reduces the α-keto acid to 2*S*-hydroxy-3-(4-methoxyphenyl) propanoic acid (MPPA) through an NADPH-dependent reaction. The C-domain then catalyzes the formation of the ester bond between the oxygen of the newly formed hydroxy function in this module 2 substrate and the carbonyl of the tethered trimethyl tyrosine residue in module 1.

Based on annotation by antiSMASH, HcaC is predicted to encode for the incorporation of an *N*-methyl valine into hectoramide B. The final module, HcaD, is also predicted to incorporate an *N*-methyl valine residue. However, it lacks a terminal thioesterase (TE) domain; instead, it possesses a domain closely related (83.5% identity, Table 2, Figure S13) to one found in the vatiamide pathway that is believed to catalyze offloading from the enzymatic pathway through terminal amidation. In the vatiamide pathway, this proposed enzymatic function is embedded in VatR, downstream of the PCP; the same domain organization is observed in the hectoramide B pathway^15^. Terminal amides resulting from an NRPS pathway have previously been found in several cyanobacterial natural products, including dragonamide A, B and E,^22^ carmabin A,^23^ and vatiamide E and F.^15^ Although the mechanism and enzymology of terminal amidation has not been studied, it is hypothesized that this motif catalyzes offloading through ammonolysis, mechanistically similar to the more typical hydrolysis of NRPS thioester linkages.

**Table 2.** Sequence identity and similarity between terminating modules HcaD and VatR.

| Amino Acid Position | Predicted function | Identity/Similarity |
| --- | --- | --- |
| 76-380 | Condensation | 23.0/43.0 |
| 555-952 | Adenylation | 88.2/95.0 |
| 1046-1267 | N-methyltransferase | 90.1/95.0 |
| 1477-1543 | Peptidyl carrier protein | 83.6/91.0 |
| 1544-1977 | Amidotransferase | 83.5/93.0 |
|  | Overall: | 72.0/86.7 |

In addition to the core domains in the *hca* gene cluster, an upstream regulatory gene, ORF 2, was identified (Table S9). This regulatory gene belongs to the *Streptomyces* Antibiotic Regulatory Protein (SARP) family, a group of transcriptional regulators commonly found in actinomycetes that regulates the biosynthesis of various antibiotic gene clusters.^24,25^ This discovery suggests that ORF 2 may play a pivotal role in controlling the production of hectoramide B. However, further analysis and investigation are necessary to fully understand its specific involvement and mechanism of action within the *hca* gene cluster. A PGAP1-like protein with hydrolase activity in ester bonds was identified in ORF1 and may also play a role in the biosynthesis of hectoramide A. It is plausible that after offloading of **1**, ORF1 catalyzes the hydrolysis of the ester bond in **1**, ultimately leading to the formation of **2**.

### Exploring the Bioactivity of Hectoramide B

To explore the potential antifungal properties of hectoramide B (**1**), a microbroth dilution method was employed. The original strain of *C. albicans* used in the co-culture experiments was no longer available, which prevented repeat testing with this strain. We therefore used *Saccharomyces cerevisiae* as a surrogate yeast strain. However, no antifungal activity was detected at a maximum test concentration of 200 μg/mL of hectoramide B. As a result, it remains uncertain whether hectoramide B (**1**) demonstrates any antifungal activity, and thus the underlying reason for its increased production in the co-culture experiment is yet unknown.

## Conclusion

Genome sequencing projects of marine cyanobacteria have revealed that they, like many other bacteria, contain a large number of orphan gene clusters that encode for cryptic natural products. Antagonistic co-culture experiments have been successful in stimulating the expression of some of these natural product BGCs^10^, and we were interested to explore this concept with our marine cyanobacterial cultures. We found hectoramide B to be prominently upregulated in the co-culture experiment with *C. albicans*. The structure of hectoramide B (**1**) was determined from a careful analysis of its spectroscopic features, comparison to the co-metabolite hectoramide A (**2**), and a bioinformatic investigation of its biosynthetic gene cluster. Although anti-yeast activity to *Saccharomyces cerevisiae* was not detected, it is still possible that hectoramide B plays a role in protecting *M. producens* JHB from antagonism by *C. albicans*. The *hca* gene cluster and its encoded protein components show several interesting features, such as an unusual motif for α-hydroxy acid incorporation, the presence of a coenzyme A ligase (CAL) in depsipeptide formation, and pathway termination through a putative ammonolysis reaction. Further biochemical interrogation of the formation of these C-terminal amides is certainly warranted. This terminal amide moiety is a popular bioisostere for improving cellular permeability of carboxylic acids;^26^ understanding how marine cyanobacteria produce this structural feature may enable development of biocatalytic methods for its creation.

## Methods

### General considerations

NMR spectra were recorded using a JEOL ECZ 500 MHz NMR spectrometer (Akishima, Tokyo, Japan). Data for NMR spectra are reported as follows: shift (δ) in ppm; s, singlet; d, doublet; t, triplet; q, quartet; m, multiplet or unresolved; brs, broad signal; *J*, coupling constant(s) in Hz. NMR spectra were analyzed using MestreNova v. 14.3.0-30573 (Mestrelab, Santiago de Compostela, Spain). Mass spectrometry data were analyzed using Xcalibur Qual Browser v. 1.4 SR1 (Thermo Fisher Scientific, Inc.). LR-LCMS data were collected on a Thermo Finnigan Surveyor Autosampler/LC-Pump-Plus/PDA-Plus with a Thermo Finnigan Advantage Max mass spectrometer. HPLC purification was carried out with a Thermo Scientific Dionex Ultimate 3000 Pump/RS/Autosampler/RS Diode Array Detector/Automated Fraction Collector using Chromeleon software. An Agilent 6230 time-of-flight mass spectrometer (TOFMS) with Jet Stream electrospray ionization source (ESI) was used for high resolution mass spectrometry (HR-MS) analysis. The Jet Stream ESI source was operated under positive ion mode with the following parameters: VCap: 3500V; fragmentor voltage: 160 V; nozzle voltage: 500 V; drying gas temperature: 325 C, sheath gas temperature: 325 C, drying gas flow rate: 7.0 L/Min; sheath gas flow rate: 10 L/Min; nebulizer pressure: 40 psi. Solvents used for extraction, purification, and LC-MS/MS analysis were purchased from Fisher Chemical. All solvents were HPLC grade except H_2_O which was purified with a Millipore Milli-Q system before use. Deuterated solvents were purchased from Cambridge Isotope Laboratories.

### Microbial strains and Culture conditions

As previously reported, *M. producens* JHB was collected from Hector’s Bay, Jamaica on August 22, 1996. The JHB culture has been continuously maintained in SWBG11 media in our laboratory at 26 ºC with a 16 h light/ 8 h dark regimen^6^ since its original collection. *Candida albicans* and *Saccharomyces cerevisiae* were stored in a -80ºC freezer in LB media with 20% glycerol and obtained from ATCC.

### Growth of *C. albicans* in SWBG11 media

LB media and SWBG11 media were prepared separately according to standard protocols.^27^ Five combinations of LB-SWBG11 media were prepared at the following ratios of LB:SWBG11: 60:40%, 40:60, 20:80, 10:90 and 100% LB media to determine optimal ratio for growth of *C. albicans* in SWBG11 media. Each media type (30 mL) was aliquoted into a 50 mL Falcon tube. One mL of *C. albicans* seed culture in LB media was added to each tube and incubated for 24 h at 37°C. Each combination was prepared in triplicate. A small amount of *C. albicans* growth was clearly observed in the lowest ratio of 10:90 LB-SWBG11 media and this condition was used for subsequent co-culture experiments.

### Co-culture of cyanobacteria with *Candida albicans*

Cyanobacteria were grown for four weeks in SWBG11 artificial seawater growth medium at 27ºC and 756 Lux in triplicate, with the following combinations: *M. producens* JHB alone and *M. producens* JHB + *C. albicans*. This light intensity was chosen because it was consistent with conditions that JHB culture was accustomed to in order to minimize changes to variables that might affect growth and productivity. Samples of *M. producens* JHB were prepared separately and added to 125 mL of SWBG11 media in a sterile, 250 mL plastic bottle. These bottles were inoculated with 5 mL of *C. albicans* from LB-SWBG11 media prepared above. Similarly, JHB in 125 mL of SWBG11 medium, were prepared as controls. The bottles were sealed and opened and gently aerated with swirling motion in a biosafety cabinet to facilitate adequate gas exchange.

### Extraction and LC-MS of Co- and Mono-cultures

After the co- and mono-cultures were incubated for 30 days, the biomass was harvested from each sample through vacuum filtration. Each culture sample was extracted four times using 2:1 dichloromethane/methanol (DCM/MeOH) by sonication for 3 min followed by soaking for 15-20 min to obtain crude extracts. Each crude extract was diluted to a concentration between 0.5-4 mg/mL (Table S14). A 1 mg portion of the extract was subjected to filtration using a C18 SPE column, dried under N_2_, and redissolved in LC-MS grade MeOH for a final concentration 1 mg/mL. The samples of the extract and a blank of MeOH were analyzed by LR-LCMS using a Phenomenex Kinetex 5 μm C18 100 A 100 × 4.60 mm column at a flow rate of 0.6 mL/min. The mobile phase was comprised of solvent A, water + 0.1% formic acid (FA) and solvent B, acetonitrile (ACN). A 32-minute method was used starting with equilibration at 30% solvent B in solvent A for 2 minutes, followed by a linear 22-minute gradient to 99% solvent B, followed by a 4-minute washout phase at 99% solvent B, and a 4-minute re-equilibration period at 30% solvent B in solvent A. Data dependent acquisition (DDA) of MS/MS spectra was performed in the positive ion mode.

### Extraction and LC-MS of hectoramide B (1)

A fresh laboratory culture sample of *M. producens* JHB was harvested through vacuum filtration resulting in 492.28 g of biomass (wet weight). It was extracted four times using 2:1 DCM/MeOH by sonication for 3 min followed by soaking for 15-20 min to afford 8.82 g of extract. A 1 mg portion of the extract was subjected to filtration using a C18 SPE column, dried under N_2_, and redissolved in LC-MS grade MeOH for a final concentration of 1 mg/mL. The samples of the extract and a blank of MeOH were subjected to LCMS and HR-MS analysis as described above.

### VLC and HPLC purification

A 4.05 g portion of the crude extract was dissolved in 4 mL of 2:1 DCM/MeOH and mixed in a 500 mL round bottom flask with 16.2 g of TLC grade silica. The mixture was dried and loaded onto a 400 mL column for vacuum liquid chromatography (VLC). Fixed 300 mL volumes of hexanes, EtOAc, and MeOH solvents were used that progressively increased in polarity: (A) 100% hexanes, (B) 10% EtOAc/hexanes, (C) 20% EtOAc/hexanes (D) 40% EtOAc/hexanes, (E) 60% EtOAc/hexanes, (F) 80% EtOAc/hexanes, (G)100% EtOAc, (H) 25% MeOH/EtOAc, and (I) 100% MeOH. Fractions H and I contained hectoramide B **(6)** and were combined and solubilized in 100 mL of EtOAc for liquid-liquid extraction with H_2_O. Three iterations of liquid-liquid extraction were performed in a separatory funnel to remove salts, giving an organic layer of 0.5253 g for combined Fractions H+I. Fractions H+I was purified by HPLC using a Thermo Scientific Dionex Ultimate 3000 Pump/RS/Autosampler/RS Diode Array Detector/Automated Fraction Collector that yielded 6 subfractions. A Phenomenex Kinetex 5 μm C18 100 Å LC 150 × 21.2 mm column was used for reverse phase separation at a flow rate of 9 mL/ min. The mobile phase consisted of solvent A, water + 0.1% formic acid (FA), and solvent B, 100% ACN. A 32-minute method was used starting with equilibration at 20% solvent B in solvent A for 5 minutes, followed by a linear 20-minute gradient to 99% solvent B, followed by a 3-minute washout phase at 99% solvent B, and a 4-minute re-equilibration period at 20% solvent B in solvent A. The third subfraction from this separation of fractions H+I contained compound **1** and was purified further by HPLC using Phenomenex Kinetex 5 μm C18 100 Å LC 100 × 4.60 mm column at a flow rate of 1 mL/ min and gradient elution as described above. This purification procedure from the monoculture of *M. producens* JHB afforded 16.5 mg of **1**.

### DNA Extraction, Nanopore Sequencing of *M. producens* JHB, and Hybrid Assembly

DNA extraction was performed using a QIAGEN Bacterial Genomic DNA Extraction Kit using the standard kit protocol. The quality of the genomic DNA (gDNA) was evaluated by Nanodrop, 1% agarose gel electrophoresis, and Qubit.

Data generation was conducted using Oxford Nanopore PromethION sequencing platform by UC Davis Genomics Core. SQK-LSK110 and FLO-PRO002 were used for library construction and data generation. All data generation was conducted using the manufacturer’s protocols. Base-calling used Guppy v5.0.7 with dna_r9.4.1_450bps_hac model. A subset of the sequencing read data was generated with Filtlong v.0.2.1^28^ with parameters Min_length=2000, keep-percent=90, and target_bases=1500000000 for read filtering.

Two long-read assembly tools (Unicycler v.0.5.0^29^ and Flye v.2.9),^30^ which can conduct assembly using only Nanopore reads or with the addition of Illumina reads, were used for this study. Flye was used for assembly using only Nanopore reads with genome-size=9m as a parameter. Unicycler was used for assembly using only Nanopore reads and in combination with Illumina reads with default parameters for hybrid assembly and long-read only assembly. Unicycler utilizes SPades, Racon and Pilon as part of the workflow. Metagenome binning was conducted using Metabat2 v2.15.^31^ The bins were annotated using CheckM v.1.2.0^32^ with taxonomy workflow, rank=phylum, and taxon=Cyanobacteria as the parameters. Bins that contained assemblies with ∼43% GC content, and total contig length of ∼9 Mbp were selected for genome polishing by Pilon.

Short reads from a previous Illumina sequencing effort were mapped to the assembly using bwa-mem2 v.0.7.17^33^ and polishing was conducted using Pilon v.1.24^34^ with default parameters. Three iterations of polishing were performed. The genome is deposited in Genbank with accession number CP017708.2.

For polished genome evaluation, BUSCO v.5.3.2^35^ was used with the cyanobacteria_odb10 database. N50, number of contigs, and genome lengths were identified using Quast v.5.0.2^36^ with default parameters. To assess BGC content, AntiSMASH v.6.0^37^ was used on the web-based platform with settings detection=relaxed, and all extra features enabled. The resulting region that contained the putative hectoramide B biosynthetic gene cluster (BGC) was downloaded as a GenBank file and investigated further using Geneious Prime v.2022.1.1.

### Sequence alignments and Phylogenetic Tree

Sequence alignments were generated using Clustal Omega v.1.2.3 on Geneious Prime software with default parameters. Methyltransferase domain and adenylation domain sequences were obtained from the NCBI and MiBIG databases.^38^ The phylogenetic tree was generated using Geneious Tree Builder with Jukes-Cantor model and default parameters. The methyltransferase (MT) domains used in the phylogenetic tree generation are listed in Table 1 of the Supporting Information. An oxygen methyltransferase from *Tistrella mobilis* KA091029-065 was used as the outgroup. Sequence alignment figures were generated by EsPript 3.0.^39^

### Structural model and alignment

The model of the hcaB adenylation domain was built using AlphaFold2^17^ with the ligand being placed by aligning to other known A domains in MOE. With the ligand placed into the predicted model of HcaB, the ligand placement was refined using a steepest descent energy minimization method followed by 10ns of low mode molecular dynamics (MD) to confirm the binding conformations of the sidechains. This final model was analyzed then in PyMol v.2.0.^40^ The model was superimposed onto other adenylation domains (A-domains) from the PDB database to obtain structural alignments (Table S12). The A-domain residues within 5 Angstroms of the binding pocket ligand in the hcaB model were evaluated as potential binding site residues by comparison with other A-domains.

### Biological Assays

Minimum inhibitory concentrations (MIC) were determined using microtiter broth dilution in Yeast Peptone Dextrose (YPD) media. Frozen spore suspensions of *Saccharomyces cerevisiae* were grown on an overnight plate culture in YPD agar. Wells were inoculated to a final concentration of 1.5 × 10^5^ cfu/mL. Plates were incubated at 30ºC for 20 hours and the MICs were defined as the lowest concentration of drug completely inhibiting visible growth. Cycloheximide and fluconazole were used as positive controls.

## ASSOCIATED CONTENT

### Supporting Information

The Supporting Information is available free of charge on the ACS Publications website.

## AUTHOR INFORMATION

### Author Contributions

The manuscript was written through contributions of all authors. All authors have given approval to the final version of the manuscript.

### Funding Sources

The research was supported by NIH grant 5R01GM107550-10 to W.H.G and L.G. K.LA. was funded by NIH/NCI T32 CA009523 and NIH T32 GM067550. C.B.N. was financially supported in part by NIH grant T32 CA009523. Research reported in this publication was supported in part by the National Center for Complementary and Integrative Health of the NIH under award number F32AT011475 to N.E.A. V,V,S. is financially supported by F32-GM145146

### Notes

Any additional relevant notes should be placed here.

## ACKNOWLEDGMENT

The sequencing was carried out by the DNA Technologies and Expression Analysis Core at the UC Davis Genome Center, supported by NIH Shared Instrumentation Grant 1s10OD010786-01. We thank Sabine Ottilie (Calibr) for providing the protocol for the yeast assay, Brendan Duggan for help with the acquisition of the 600 MHz NMR data sets and Dr. Yongxuan Su for help with acquisition of the HRMS data set. We thank the Dickinson Foundation for purchase of the JEOL ECZ 500 MHz NMR Spectrometer.

## ABBREVIATIONS

A: adenylation
ACN: acetonitrile
AMT: amidotransferase
BGC: biosynthetic gene cluster
C: condensation
CAL: co-enzyme A ligase
COSY: correlated spectroscopy
DCM: dichloromethane
EtOAc: ethyl acetate
FA: formic acid
gDNA: genomic DNA
GNPS: global natural products social molecular networking
HMBC: heteronuclear multiple bond correlation
HPLC: high performance liquid chromatography
HSQC: heteronuclear single quantum coherence
JHB: *Moorena producens* JHB
KR: ketoreductase
LC-MS: liquid chromatography mass spectrometry
MeOH: methanol
MIC: minimum inhibitory concentration
Mpopa: 3-(4-methoxyphenyl)-2-oxopropanoic acid
Mppa: 2-hydroxy-3-(4-methoxyphenyl) propanoic acid
MT: methyltransferase
NMR: nuclear magnetic resonance
NMT: nitrogen-methyltransferase
NRP: non-ribosomal peptide
NRPS: non-ribosomal peptide synthetase
OMT: oxygen-methyltransferase
PCP: peptidyl-carrier protein
SARP: streptomyces antibiotic regulatory protein
TE: thioesterase
VLC: vacuum liquid chromatography.

## Notes

### Competing Interest Statement

The authors have declared no competing interest.

