## Supplementary Information for "Structure and Biosynthesis of Hectoramide B, a Linear Depsipeptide from the Marine Cyanobacterium *Moorena producens* JHB Discovered via Co-culture with *Candida albicans*"

Table of Contents:

**Spectroscopic Data for Hectoramide B (1)**

**S1**	NMR data summary for hectoramide B (**1**).

**S2**	^1^H-NMR spectrum (600 MHz, MeOH-*d*_4_) of hectoramide B (**1**).

**S3**	^1^H-NMR spectrum of hectoramide B (**1**) with solvent suppression.

**S4**	HSQC-NMR spectrum (600 MHz, MeOH-*d*_4_) of hectoramide B (**1**).

**S5**	HMBC-NMR spectrum (600 MHz, MeOH-*d*_4_) of hectoramide B (**1**).

**S6**	COSY-NMR spectrum (600 MHz, MeOH-*d*_4_) of hectoramide B (**1**).

**S7**	SMART-NMR analysis of HSQC spectrum of hectoramide B (**1**)

**Biosynthetic Gene Cluster Analysis of Hectoramide B (1)**

**S8**	Retrobiosynthetic scheme for hectoramide B (**1**).

**S9**	Deduced functions of proteins in *hca* biosynthetic gene cluster.

**S10**	Phylogenetic tree of oxygen- and nitrogen-methyltransferase (MT) domains from cyanobacteria

**S11**	Methyltransferase domains from cyanobacteria used in the phylogenetic tree shown in Figure S10.

**S12**	Adenylation domains used for sequence and structural alignment with HcaB-A domain.

**S13**	Sequence alignment of hcaD and vatR terminating module.

**LC-MS Analysis of Co- and Mono-cultures**

**S14**	Summary table of crude extracts obtained from co- and mono-cultures.

**S15**	Analysis of Relative Abundance of Secondary Metabolites in Co- and Mono-culture

**S16**	HR-MS spectra of hectoramide B (**1**)

**S1.** NMR Data Summary for Hectoramide B (**1**).

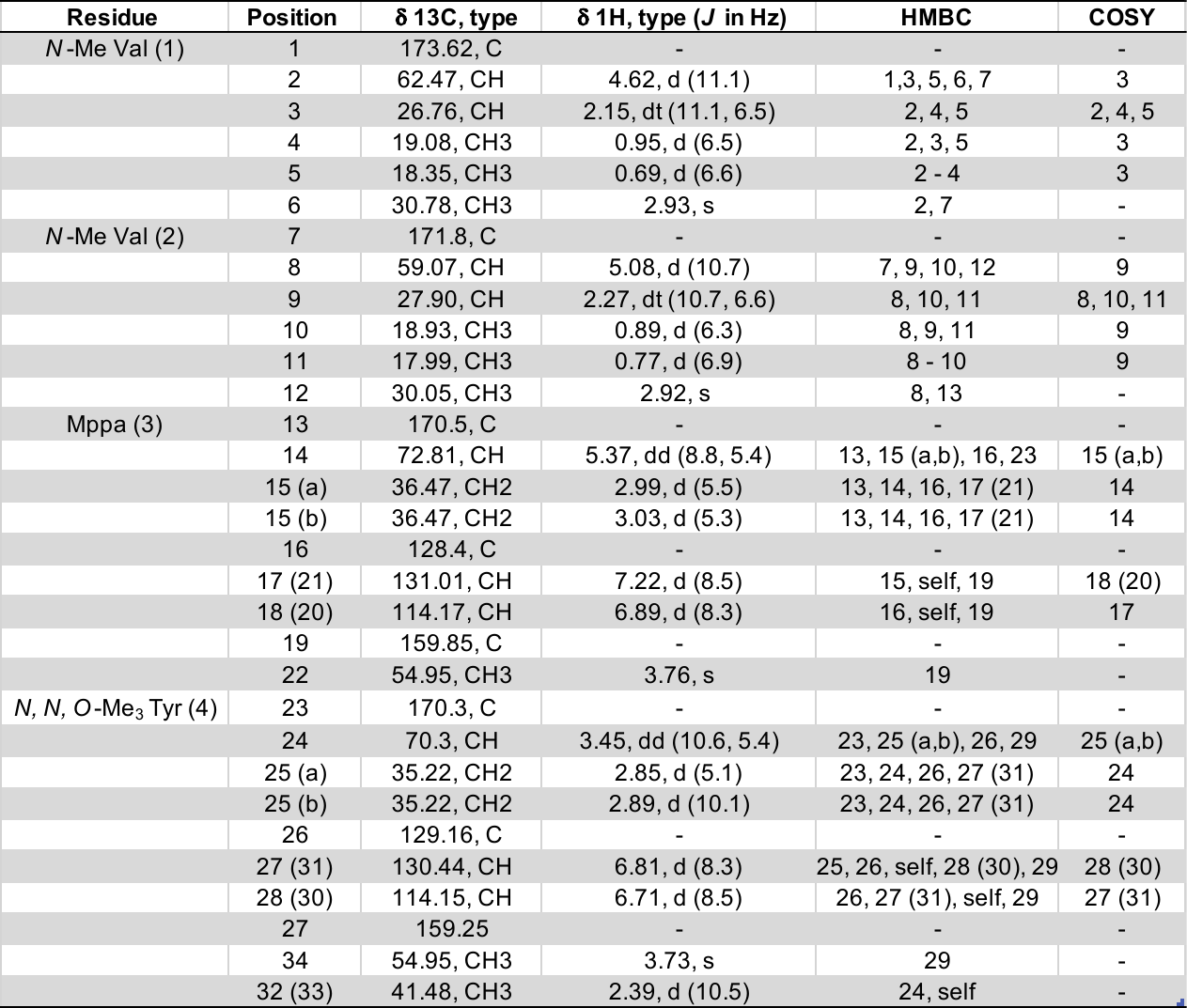

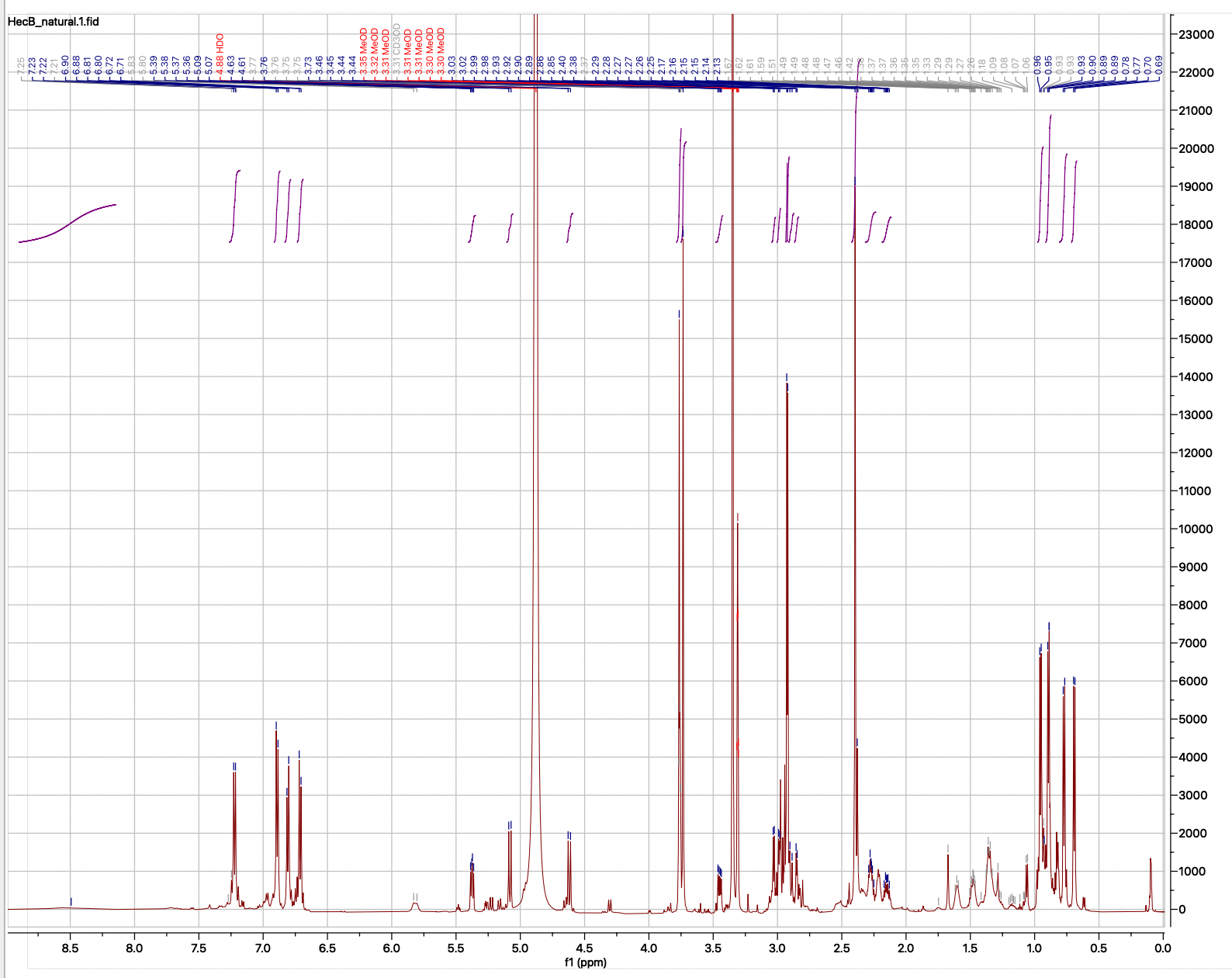

**S2.** ^1^H-NMR spectrum (600 MHz, MeOH-*d*_4_) of hectoramide B (**1**).

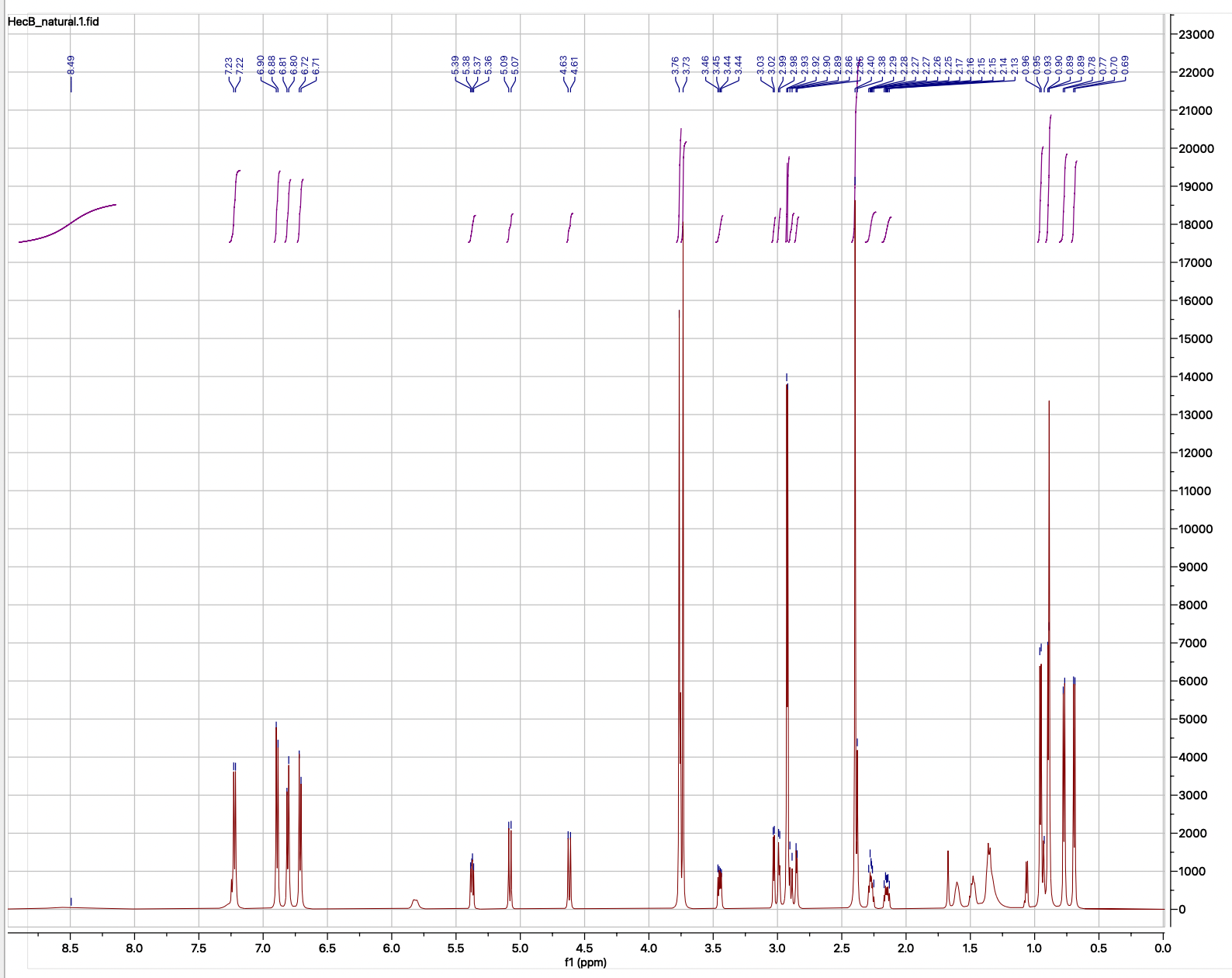

**S3.** ^1^H-NMR spectrum (600 MHz, MeOH-*d*_4_) of hectoramide B (**1**) with solvent suppression.

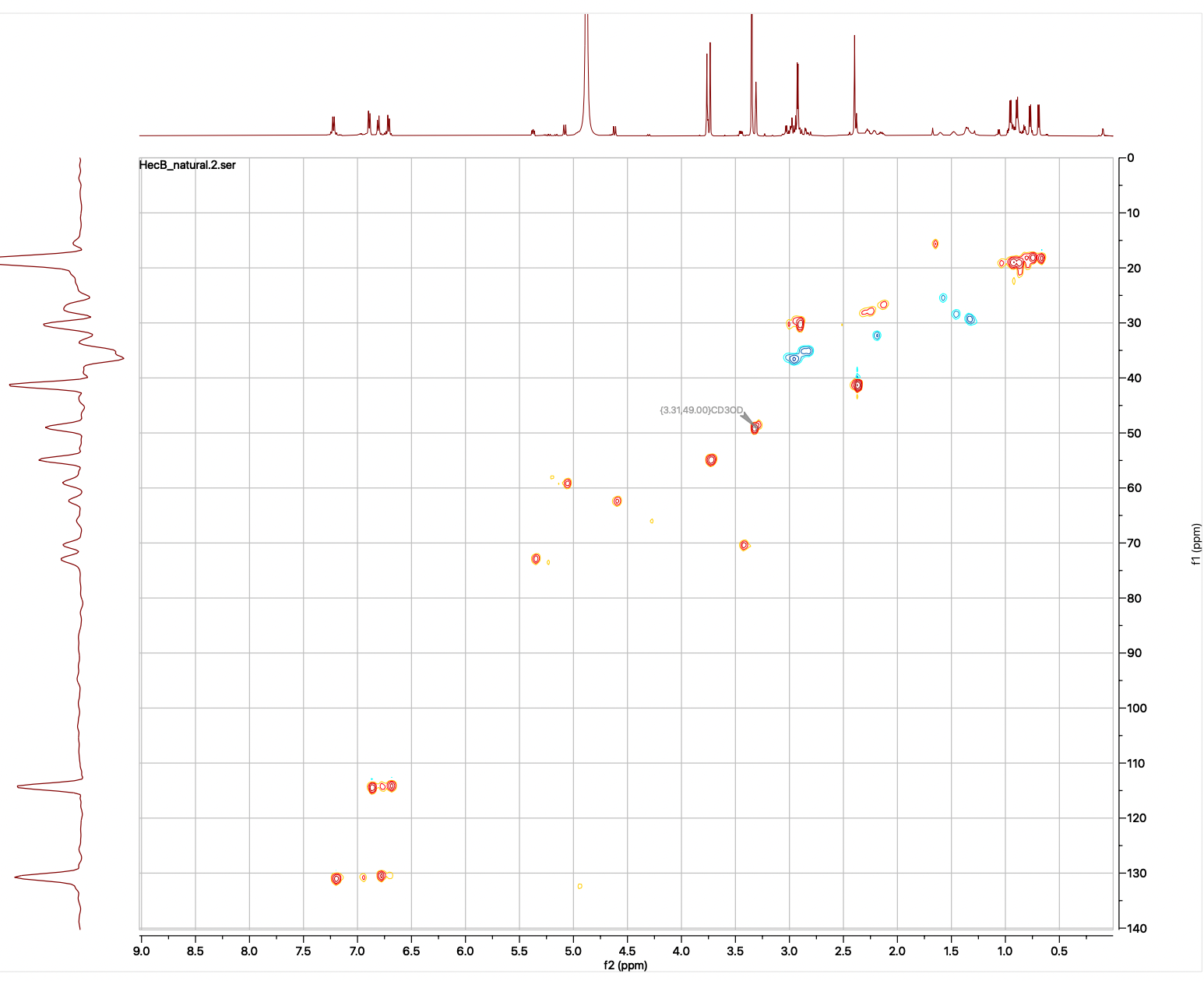

**S4.** HSQC-NMR spectrum (600 MHz, MeOH-*d*_4_) of hectoramide B (**1**).

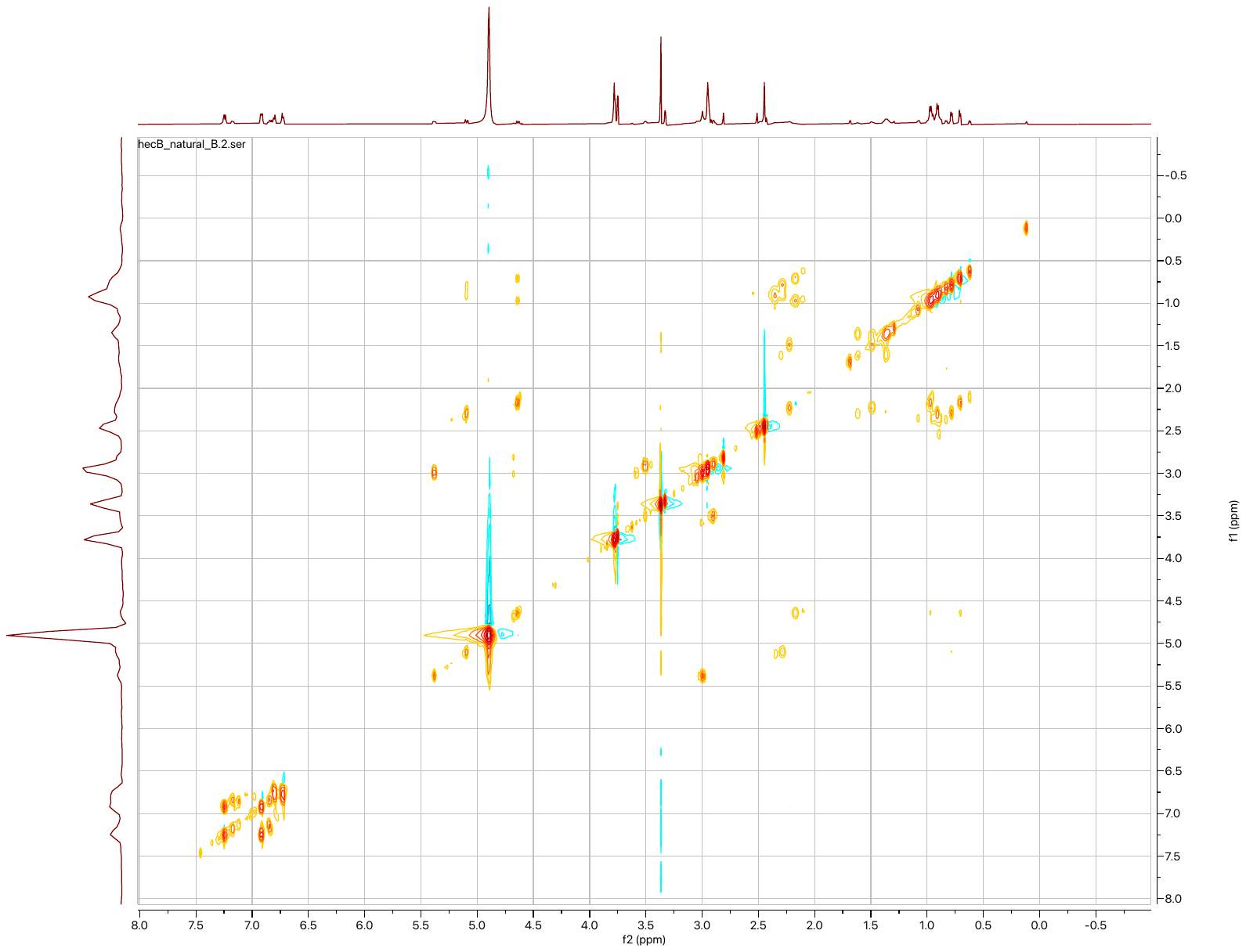

**S5.** COSY-NMR spectrum (600 MHz, MeOH-*d*_4_) of hectoramide B (**1**).

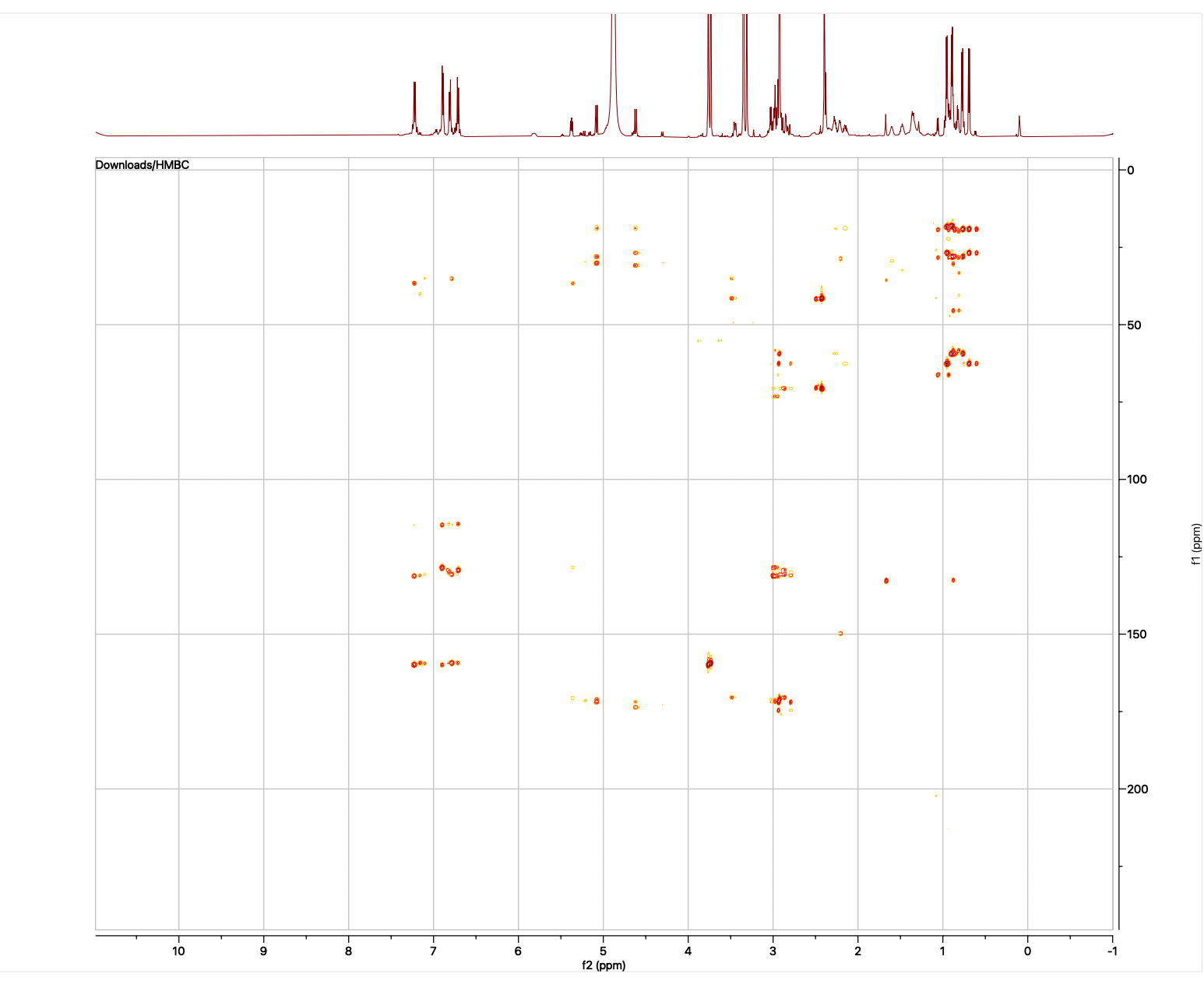

**S6.** HMBC-NMR spectrum (600 MHz, MeOH-*d*_4_) of hectoramide B (**1**).

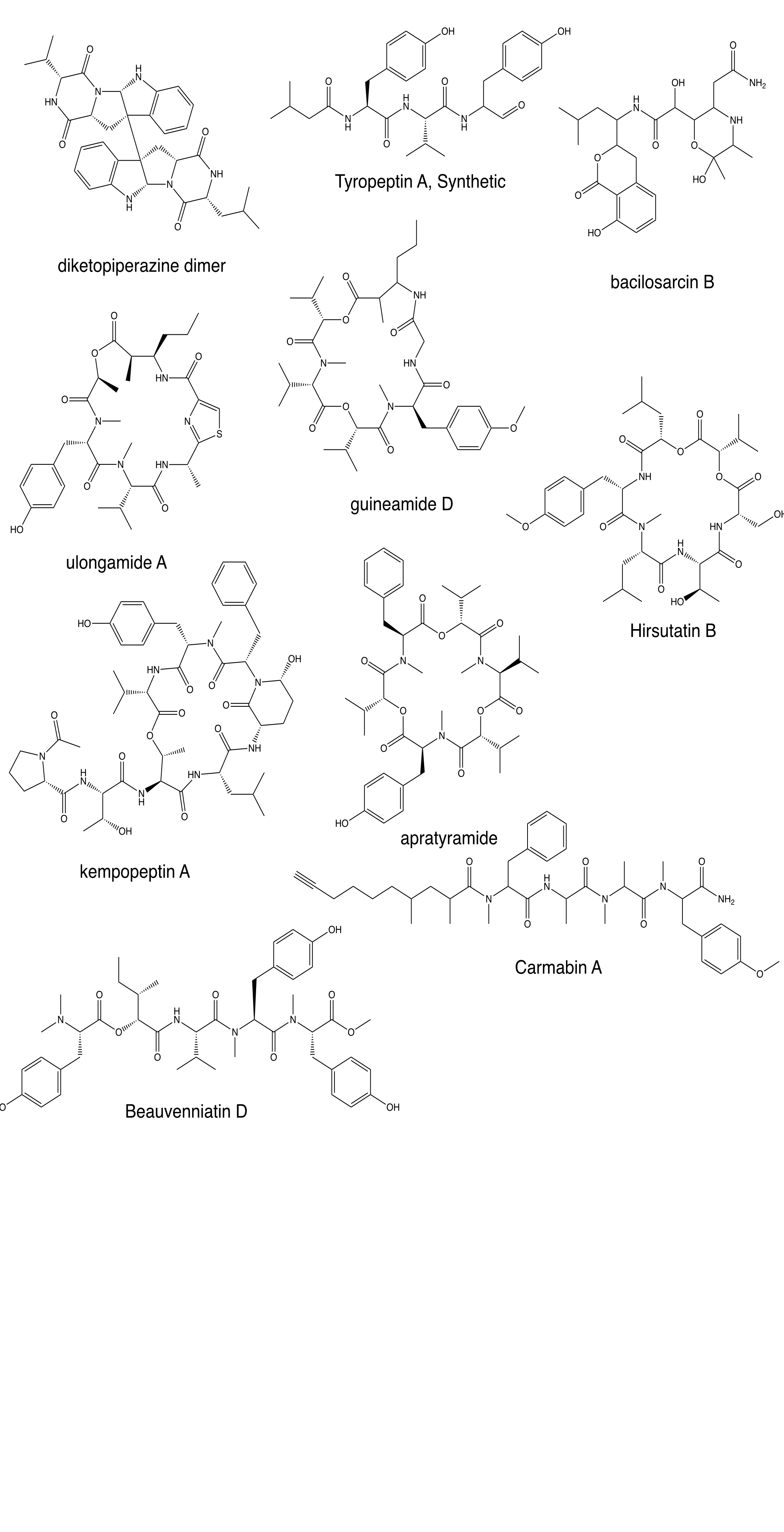

**S7**. Structures related to hectoramide B based on the SMART-NMR analysis of HSQC spectrum of hectoramide B (**1**)

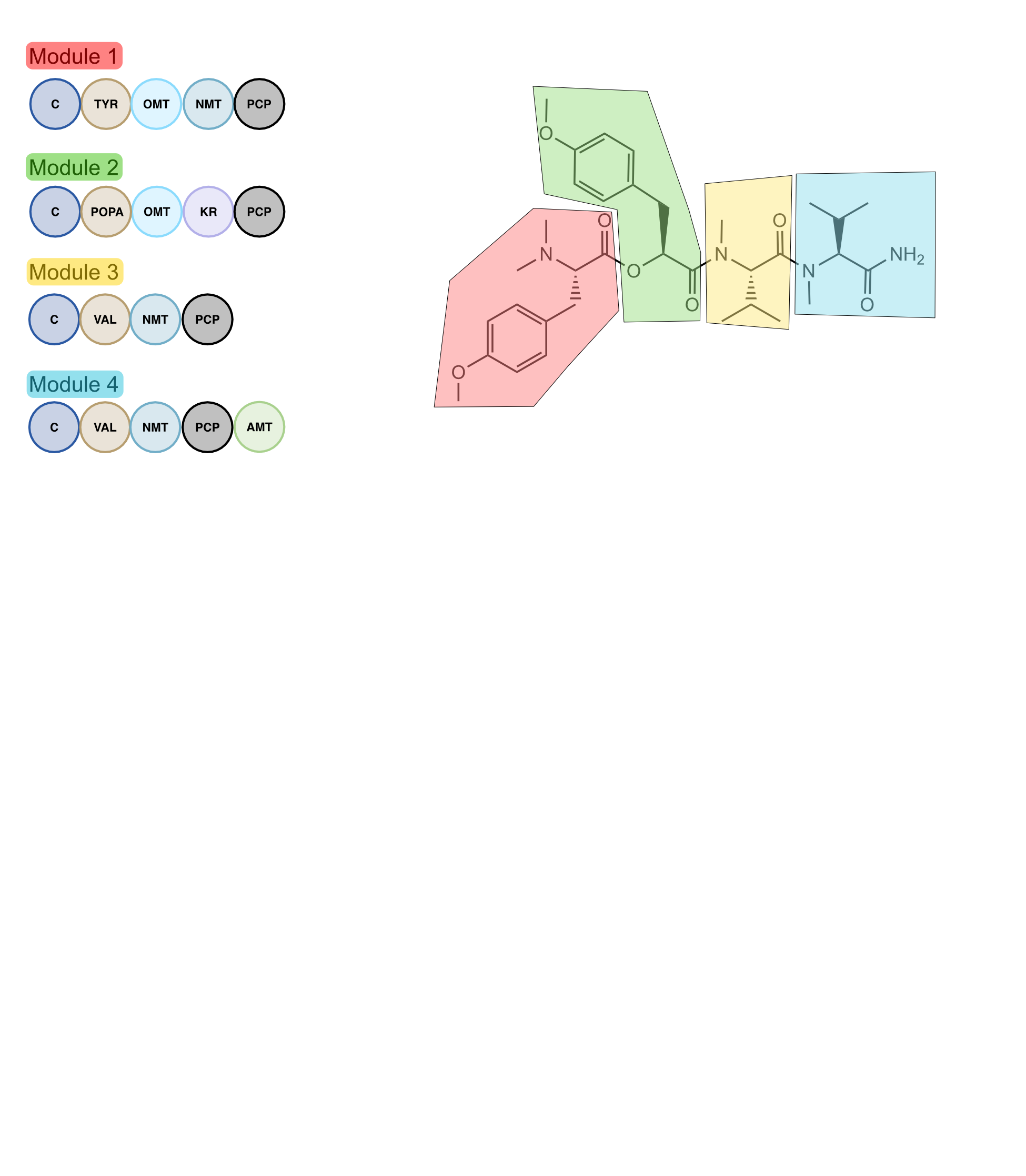

**S8.** Retrobiosynthetic scheme of hectoramide B (**1**) with the predicted modular organization of the hectoramide B biosynthetic gene cluster. Circles represent enzymatic domains. C: condensation domain, TYR: adenylation domain for tyrosine, POPA: adenylation domain for 3-(4-hydroxyphenyl)-2-oxopropanoic acid, VAL: adenylation domain for valine, KR: ketoreductase domain, NMT: Nitrogen-methyltransferase domain, OMT: oxygen-methyltransferase domain, PCP: peptidyl-carrier protein, AMT: amidotransferase

**S9.** Deduced functions of the proteins in the *hca* biosynthetic gene cluster.

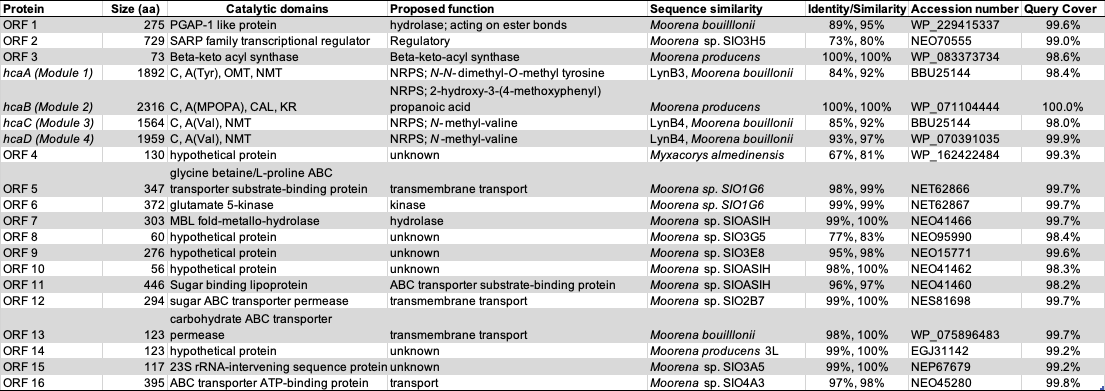

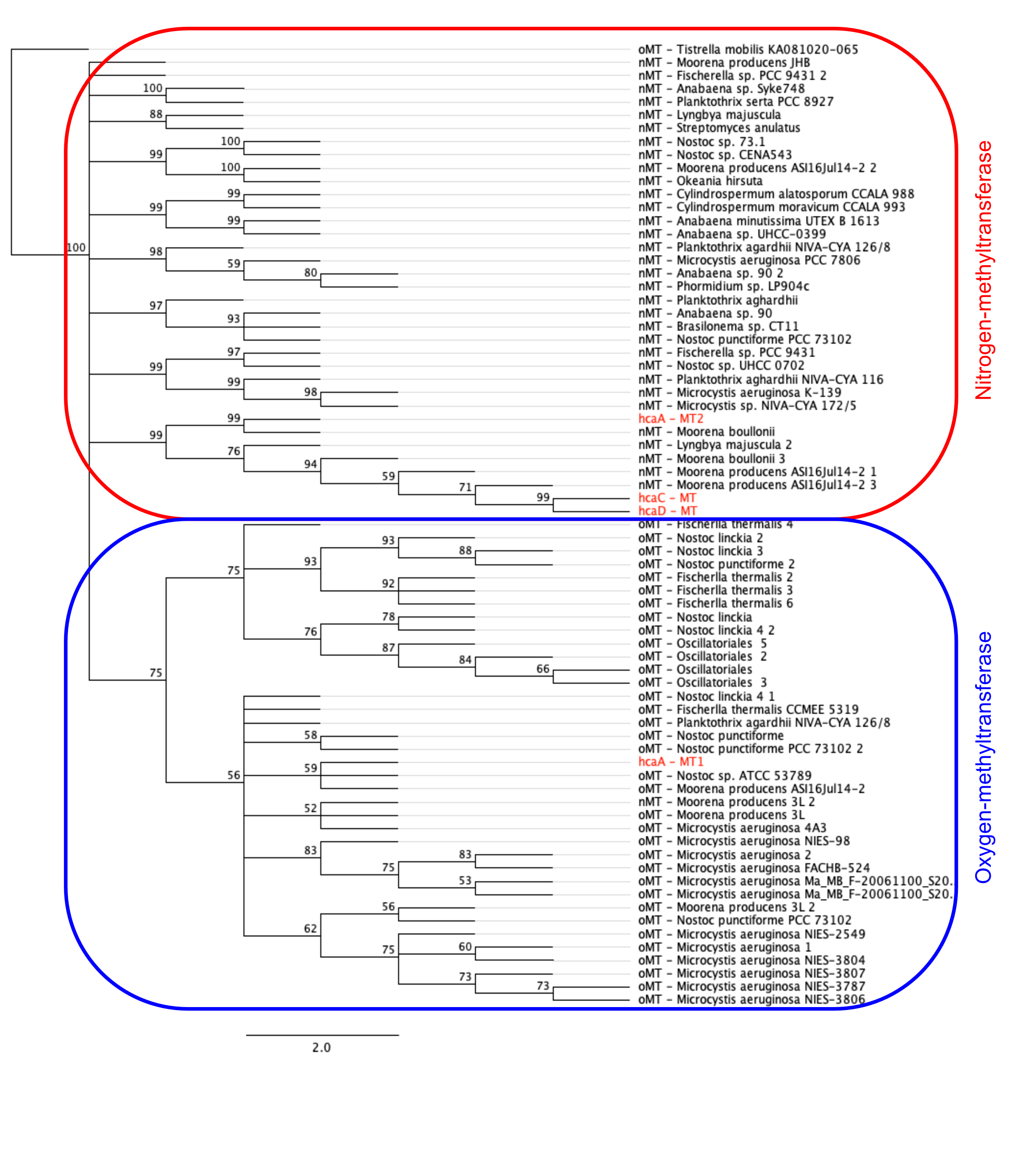

**S10**. Phylogenetic tree of oxygen- and nitrogen-methyltransferase (MT) domains from cyanobacteria reveals the specificity of the two MT domains encoded by *hcaA*. OMT domain clades are outlined in blue. NMT domain clades are outlined in red.

**S11.** Methyltransferase domains from cyanobacteria used to build the phylogenetic tree shown in Figure S10.

Sequences were prepared from NCBI database and MiBIG database.

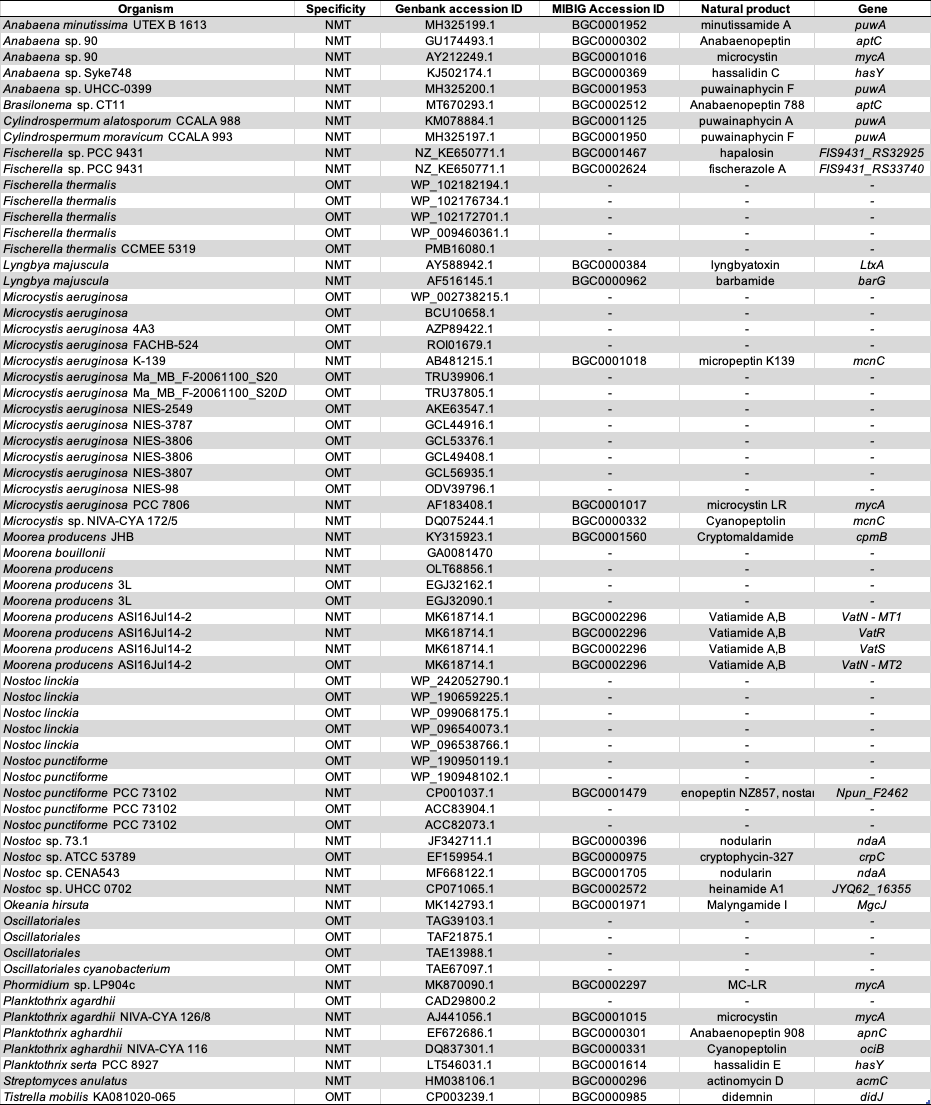

**S12.** Adenylation domains used for sequence and structural alignment with HcaB adenylation domain.

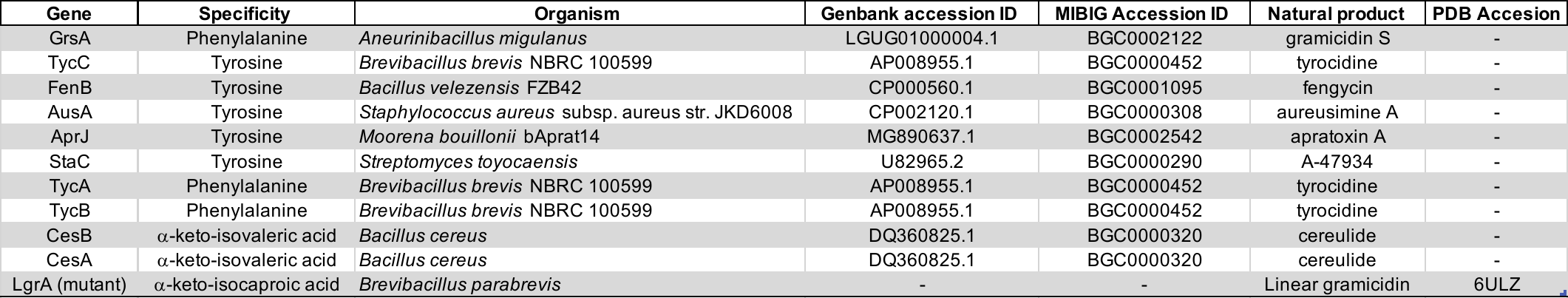

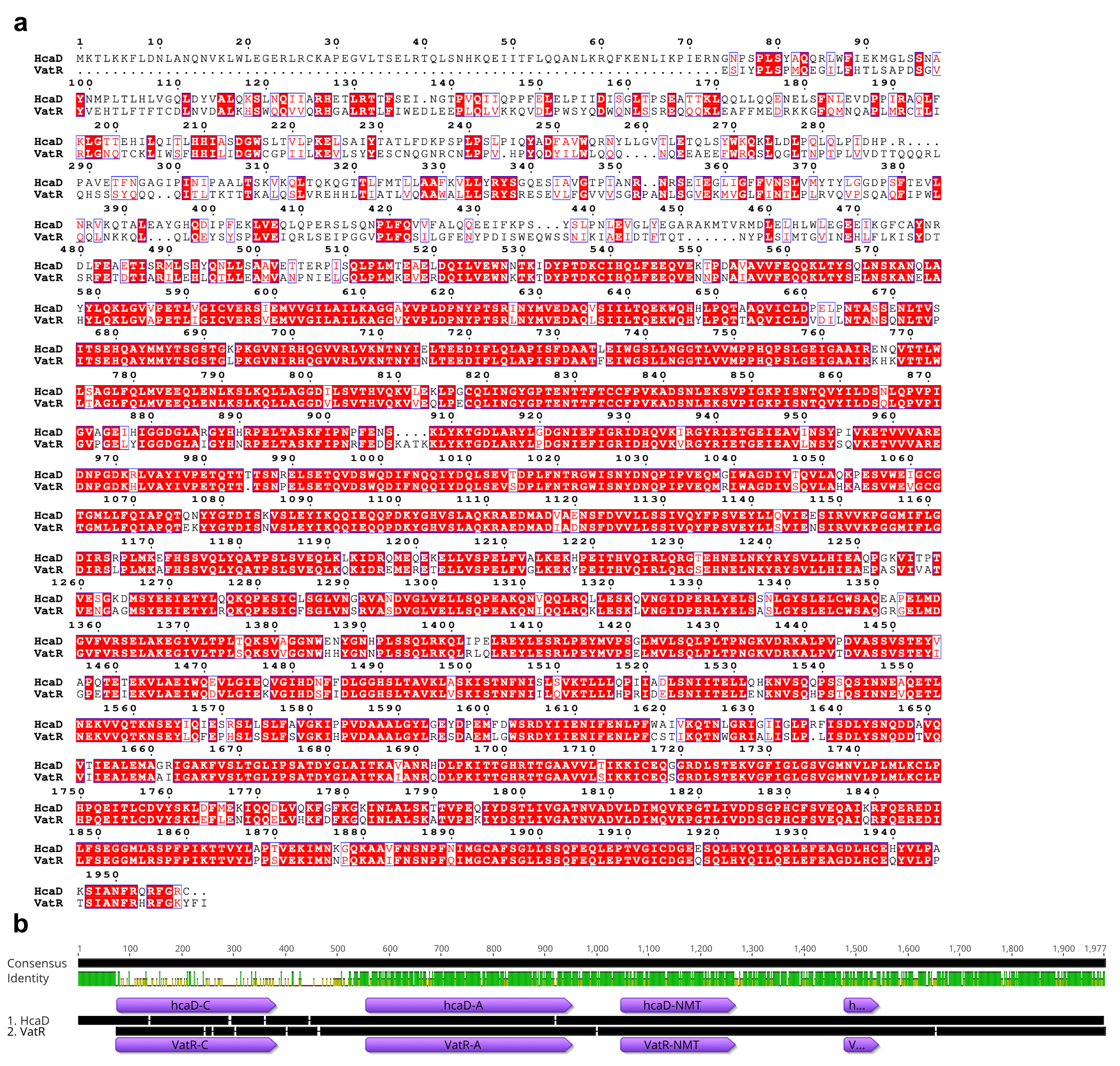

**S13.** Sequence alignment of hcaD and vatR terminating module. (a) Both pathways are predicted to encode enzymes that incorporate an *N*-methyl valinamide terminus. While these two pathways show 72% identity when aligning all 1885 amino acids, they are 93.0% identical for the 415 residues of the proposed terminating ammonolysis domain. Identical residues are highlighted in red, similar residues are outlined in blue and presented in red text. (b) Sequence alignment in gene graphic format

**S14.** Summary table of crude extracts obtained from co- and mono-cultures. JHB: *Moorena producens* JHB; CA: *Candida albicans*

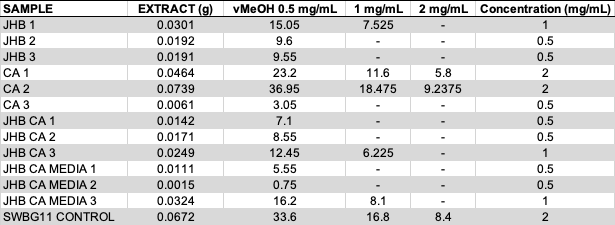

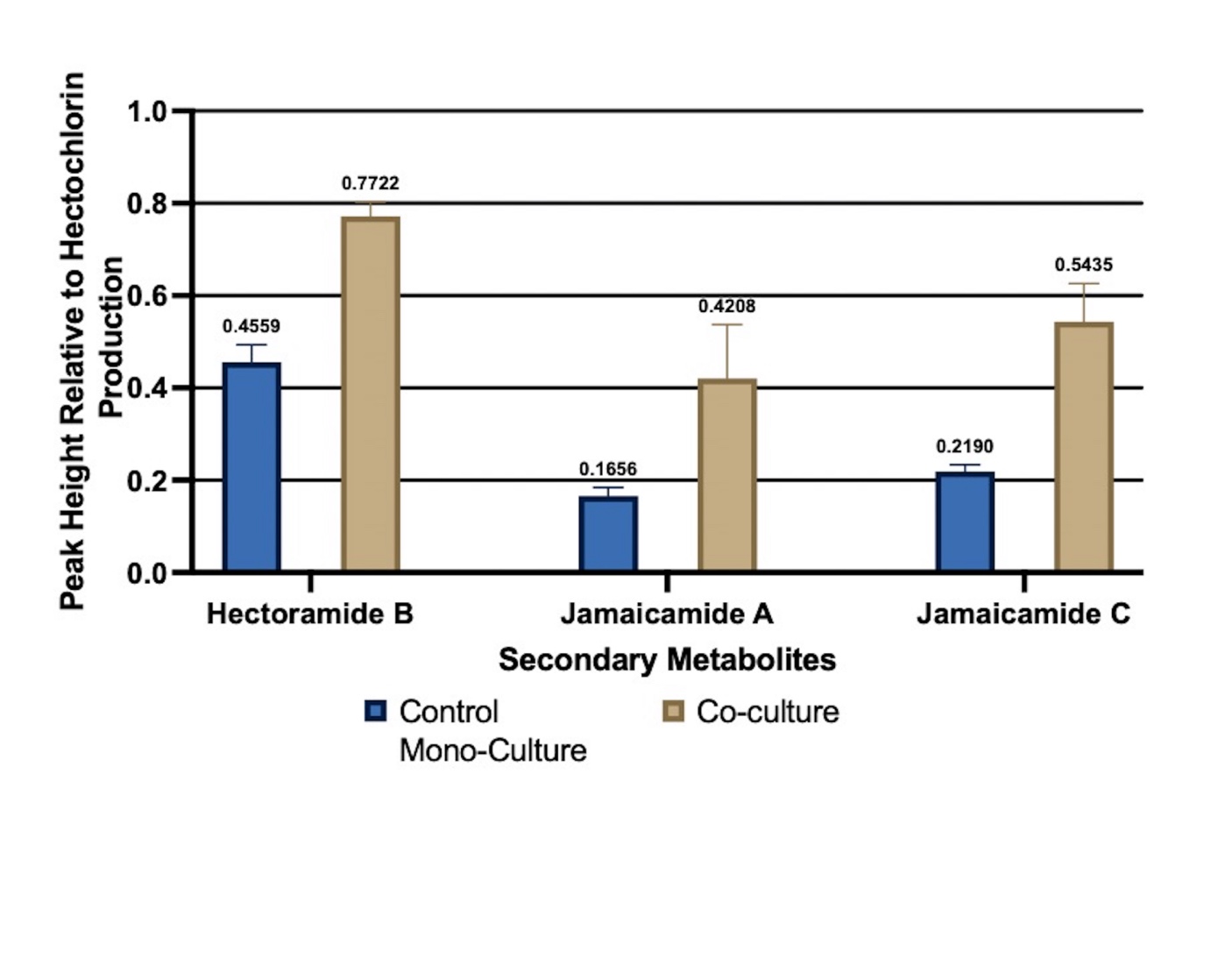

**S15. Analysis of Relative Abundance of Secondary Metabolites in Co- and Mono-cultures.** Significant increase in the production of hectoramide B as well as two known, bioactive jamaicamides when normalized to the hectochlorin content in each sample. Hectoramide A (**2**) was not detected in these experiments.

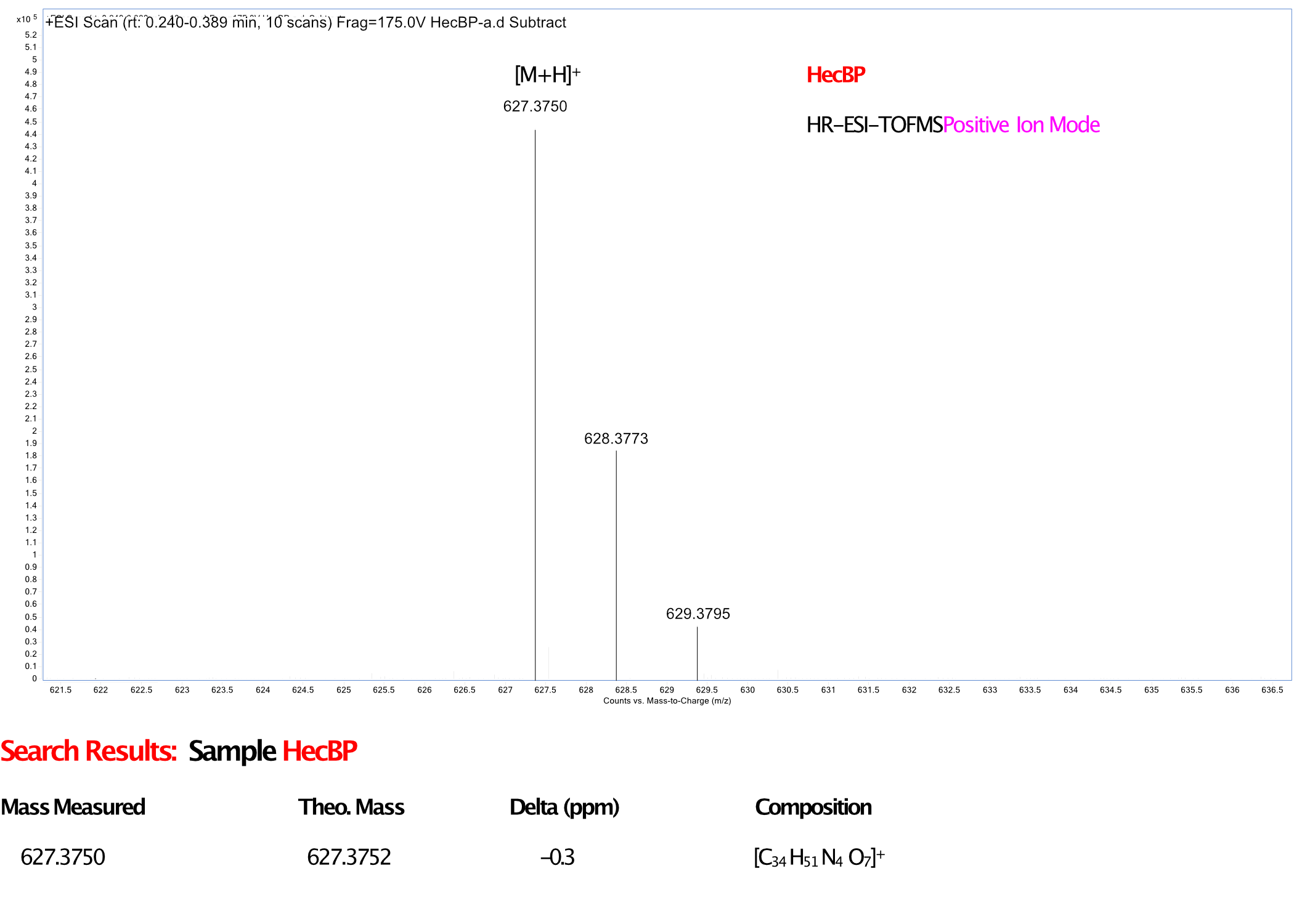

**S16**. **HRMS spectra of hectoramide B (1)**
